# Electrospun polyester-urethane scaffold preserves mechanical properties and exhibits strain stiffening during *in situ* tissue ingrowth and degradation

**DOI:** 10.1101/783522

**Authors:** Hugo Krynauw, Rodaina Omar, Josepha Koehne, Georges Limbert, Neil H Davies, Deon Bezuidenhout, Thomas Franz

**Affiliations:** Division of Biomedical Engineering, Department of Human Biology, University of Cape Town, Observatory, South Africa; Cardiovascular Research Unit, Chris Barnard Division of Cardiothoracic Surgery, University of Cape Town, Observatory, South Africa; National Centre for Advanced Tribology & Bioengineering Science Research Group, Department of Mechanical Engineering Sciences, Faculty of Engineering and Physical Sciences, University of Southampton, Southampton, UK

**Keywords:** Electrospinning, elastic modulus, mechanical properties, soft tissue regeneration, degradation

## Abstract

Consistent mechanical performance from implantation through healing and scaffold degradation is highly desired for tissue-regenerative scaffolds, e.g. when used for vascular grafts. The aim of this study was the paired *in vivo* mechanical assessment of biostable and fast degrading electrospun polyester-urethane scaffolds to isolate the effects of material degradation and tissue formation after implantation. Biostable and degradable polyester-urethane scaffolds with substantial fibre alignment were manufactured by electrospinning. Scaffold samples were implanted paired in subcutaneous position in rats for 7, 14 and 28 days. Morphology, mechanical properties and tissue ingrowth of the scaffolds were assessed before implantation and after retrieval. Tissue ingrowth after 28 days was 83 ± 10% in the biostable scaffold and 77 ± 4% in the degradable scaffold. For the biostable scaffold, the elastic modulus at 12% strain increased significantly between 7 and 14 days and decreased significantly thereafter in fibre but not in cross-fibre direction. The degradable scaffold exhibited a significant increase in the elastic modulus at 12% strain from 7 to 14 days after which it did not decrease but remained at the same magnitude, both in fibre and in cross-fibre direction. Considering that the degradable scaffold loses its material strength predominantly during the first 14 days of hydrolytic degradation (as observed in our previous *in vitro* study), the consistency of the elastic modulus of the degradable scaffold after 14 days is an indication that the regenerated tissue construct retains it mechanical properties.

## 1. Introduction

Regenerative medicine has emerged as one of the most dynamic drivers in the development of advanced engineered biomaterial solutions for tissue engineering applications [1, 2]. One crucial element in regenerative medicine are scaffolds that facilitate and guide the regeneration of biological tissues according to the biophysical requirements of the specific application.

The ideal scaffold facilitates tissue regeneration such that the new tissue constructs mimics biologically and mechanically the healthy host tissue [2]. In cardiovascular regenerative therapies, one of the major challenges remains to be small-diameter vascular grafts [3]. For these grafts, a key factor for the long term success is porosity [3]. One method of introducing porosity in scaffolds for the vascular tissue engineering is electrospinning [4-9]. Process parameters allow controlling the degree of alignment of the fibres of the resulting polymeric network [10, 11]. Fibre alignment strongly determines the mechanical properties of the scaffold [12]. Random fibre alignment leads to mechanical isotropy of the scaffold, which exhibits similar properties in different directions. In contrast, the predominant alignment of fibres in one direction introduces mechanical anisotropy [11]. Electro-spun scaffolds with a high degree of fibre alignment exhibit the highest elastic modulus, i.e. stiffness, in direction of the predominant fibre alignment and the lowest elastic modulus perpendicular to the fibre alignment. Mechanical anisotropy can offer a useful tool to tailor the structural properties of the scaffold but also adds complexity to the design process. The latter needs to be addressed adequately, in particular for biodegradable scaffolds and considering that tissue regenerating in the scaffold will contribute to structural and mechanical properties of the regenerating tissue construct.

Scaffold degradation and tissue incorporation are transient processes, and gradual change of the mechanical properties of the scaffold-tissue construct is expected until degradation and tissue ingrowth are complete. Biomechanical properties of scaffolds were predominantly obtained prior to tissue culturing or implantation [13, 14, 7, 4, 15, 16] whereas the changes in mechanical properties due to *in situ* tissue ingrowth after implantation in fast-degrading scaffolds have received very little attention.

In previous studies, we separately quantified the effect of *in vitro* degradation [17] and *in situ* tissue ingrowth in a biostable scaffold [18] on the scaffolds’ mechanical properties. The present study aimed at investigating the combined effects of concurrent *in situ* tissue ingrowth and scaffold degradation on the mechanical properties of a fast-degrading electrospun polyester-urethane scaffold. Matched implants of the polyester-urethane scaffold with a fast-degrading and a non-degrading formulation, respectively, served as internal controls.

## 2. Materials and Methods

### 2.1 Material

DegraPol® (ab medica S.p.A, Cerro Maggiore, MI, Italy), a biodegradable polyester-urethane that consists of poly(ε-caprolactone-co-glycolide)-diol soft segments and poly(3-(R-hydroxybutirrate)-co-(ε-caprolactone))-diol hard segments was used. Both polymer segments are biodegradable and their degradation products are non-toxic [19]. The mechanical properties of the polymer can be modulated by adjusting the ratio of the hard and soft segments. Changing the ratio of ε-caprolactone to glycolide affects the degradation characteristics. DegraPol® DP0 and DP30 have ε-caprolactone-to-glycolide ratio of 100:0 and 70:30 respectively, and both have a hard-to-soft segment ratio of 40:60 (unpublished data).

### 2.2 Electro-spinning and sample preparation

DegraPol® DP0 and DP30 were each dissolved in chloroform at room temperature with subsequent sonication in distilled water at 37°C for 90 min to obtain polymer solutions of 24% by weight concentration. The DegraPol® solution was electro-spun from a hypodermic needle (SE400B syringe pump, Fresenius, Bad Homburg, Germany) onto a rotating and bi-directionally translating tubular target (hypodermic tubing, Small Parts, Loganport, IN, USA) using a custom-made rig. The spinning parameters were: solution flow rate of 4.8 mL/h, target outer diameter of 25 mm, target rotating speed of 9600 rpm, target translational speed of 2.6 mm/min, electrostatic field of 15 kV, and source-target distance of 250 mm. After completion of the spinning process, the electro-spun structure on the target hypotube was submersed in ethanol for 5 minutes, removed from the mandrel and dried under vacuum (Townson & Mercer Ltd, Stretford, England; room temperature, 90 min) and trimmed on either end to discard regions with decreasing wall thickness. Tubular scaffolds were cut into rectangular samples with the longer edge aligned with either the circumferential (C) or axial (A) direction of the tube. Samples underwent standard ethylene oxide sterilisation (55°C, 60% relative humidity, 12 h) and were subsequently placed in vacuum for 6 h to allow removal of ethylene oxide.

### 2.3 Physical characterisation of scaffold before implantation

Physical characterisation included microscopic analysis of fibre diameter and alignment, measurement of scaffold wall thickness and porosity as described previously [18].

#### Fibre diameter and alignment

Scanning electron microscope (SEM) images were obtained (Nova NanoSEM 230, FEI, Hillsboro, OR, US) of samples after sputter-coating with gold (Polaron SC7640, Quorum Technologies, East Grinstead, UK). The fibre diameter was measured on x750 SEM micrographs with Scion Image (Scion Corporation, Frederick, USA) (n = 3 samples; 4 images per sample, i.e. one image from two different locations of each sample surface; ten measurements per image). Fibre alignment was quantified on x100 SEM images using Fiji [20], by means of Fourier component analysis with the Directionality plug-in (written by JeanYves Tinevez) as described by Woolley et al. [21]. The analysis provides a dispersion factor as a measure of alignment of the fibres with smaller dispersion values indicating higher degree of alignment.

#### Sample dimensions

Width, length and thickness of scaffold samples were measured on macroscopic images obtained with a Leica DFC280 stereo microscope using Leica IM500 imaging software (Leica Microsystems GmbH, Wetzlar, Germany).

#### Scaffold porosity

Scaffold porosity, P, was determined by hydrostatic weighing typically employed for density determination. The porosity of fibrous networks is described as volumetric ratio

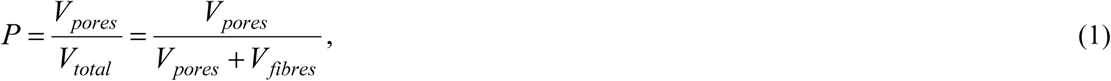

and when expressing volumes as masses in the form of

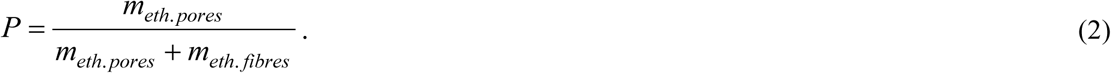

Introducing measurable quantities *m*_*wet*_, *m*_*dry*_ and *m*_*submerged*_ provides

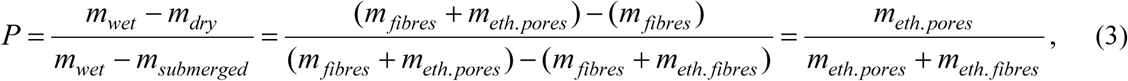

with *m*_*dry*_ as the mass of the dry scaffold, *m*_*submerged*_ as the mass of the scaffold submerged in ethanol, and *m*_*wet*_ as the scaffold mass after removal from the ethanol while ethanol is retained in the pores. The porosity was measured with n = 1 per spun scaffold.

### 2.4 Molecular weight

Weight-average molar weight, M_W_, during degradation was determined by gel permeation chromatography (GPC) using tuneable absorbance detector (Waters 486, set to 260 nm), high-performance liquid chromatography pump (Waters 510), differential refractometer (Waters 410) and auto sampler (Spectra Series AS100, Thermo Separations). Five columns (Styragels HR1 through HR6, Waters) and a pre-column filter were used at 30°C. Polystyrene standards were used for calibration. Samples were dissolved in tetrahydrofuran (1 mg/ml, 37°C, sonication for 20 min). A sample volume of 180 µl was injected with a flow rate of 1.06 ml/min.

### 2.5 Study design

This study used a subcutaneous rat model with implant durations of 7, 14 and 28 days and n = 10 animals per each time point. Each animal received six implants, namely thee biostable DP0 scaffold samples and three biodegradable DP30 scaffold samples.

### 2.6 Implantation procedure

All animal experiments were authorised by the Faculty of Health Sciences Research Ethics Committee of the University of Cape Town and were performed in accordance with the National Institutes of Health (NIH, Bethesda, MD) guidelines.

Male Wistar albino rats with body mass 200-250 g were used. Anaesthesia was induced by placing the animal in an inhalation chamber with an air flow of 5% isoflurane for 5 min. The animals were shaved and sterilized with iodine in the area of the incision. Anaesthesia was maintained by nose cone delivery of 1.5% isoflurane at an oxygen output of 1.5 L/min at 1 bar and 21°C. The body temperature of the animals was maintained throughout the surgical procedure by placing the animal on a custom-made heating pad at 37°C.

Six longitudinal incisions of 1 cm (three on either side of the dorsal midline) were made and subcutaneous pockets of 2 cm depth bluntly dissected. Scaffold samples were placed within the pockets and incisions closed with silk 4/0 (Ethicon Inc, Somerville, NJ) interrupted sutures in a subcuticular fashion.

### 2.7 Implant retrieval

Anaesthesia was induced by placing the animal in an inhalation chamber with an air flow of 5% isoflurane for 5 min. Rats were euthanized by inhalation of 5% halothane in air followed by a cardiac injection of 1 ml saturated KCl solution. The samples with surrounding tissue were excised. Samples for mechanical testing were submerged in phosphate buffered saline (PBS) solution and stored at 4°C. Samples for histological analysis underwent tissue fixation in 10% formalin (Sigma Aldrich, Steinheim, Germany) for 48 hours and transferred into 70% ethanol solution for storage.

### 2.8 Mechanical testing

Prior to testing, the thickness, length and width of retrieved samples were determined with a calliper. Tensile testing was performed within 12 hours of implant retrieval with samples submerged in PBS at 37°C (Instron 5544, 10 N load cell; Instron, Norwood, USA). Samples were fixed using custom-made clamps resulting in a gauge length of approximately 10 mm. The direction of tensile load represented either circumferential (C samples) or axial direction (A samples) of the original tubular scaffold.

Pre-implant samples (T = 0 days) underwent a loading protocol comprising either (1) one initial extension to 12% strain, five cycles between 12% and 8% strain, and a final extension to failure or maximum force of 7.4 N (all at strain rate of 9.6 %/min); or (2) one single extension to 16% strain at 9.6 %/min. Retrieved implant samples were not subjected to loading cycling but underwent one single extension only (to 16% strain at 9.6 %/min) to avoid inducing structural damage in the degraded samples at the latter stages of the study. Stress-strain data obtained for pre-implant scaffold samples with and without load-cycling was used to derive a scaling function S_C_(ε) = σ_Cycled_(ε)/σ_Noncycled_(ε) that quantifies the mechanical effect of the load-cycling. Stress-strain data obtained without load cycling on retrieved scaffold samples were adjusted using S_C_ for each test and average data calculated. Characteristic mechanical measures were derived from these data, namely stress σ_12%_ at 12% strain and σ_16%_ at 16% strain, and elastic modulus E_6%_ at 6% strain and E_12%_ at 12% strain. (Note: Stress and strain always refers to nominal stress and Cauchy strain, respectively.)

### 2.9 Histology, light microscopy and assessment of tissue ingrowth

Fixed samples underwent tissue processing and paraffin embedding. Sections of 3 μm thickness were prepared from mid sample regions and stained with haematoxylin and eosin (H&E) for nuclei and cytoplasm. Microscopic images were acquired with a Nikon Eclipse 90i microscope with digital camera DXM-1200C (Nikon Corporation, Tokyo, Japan).

Tissue ingrowth was quantified by morphometric image analysis classifying areas as open space or tissue (VIS Visiopharm Integrator System, Visiopharm, Hørsholm, Denmark). A region of interest (ROI) in the image was segmented in tissue (nuclei, cytoplasm and extracellular matrix) and open space with an untrained k-means clustering technique. The algorithm classified scaffold fibres as open space since the polymer did not stain. This was corrected by adjusting for scaffold porosity.

### 2.10 Statistical analysis

For quantitative data, one-way ANOVA was performed when more than two groups were compared, using Tukey HSD post-hoc analysis with p < 0.05 indicating statistical significance. Data are provided as mean values ± standard deviation.

## 3. Results

SEM micrographs of the electrospun DegraPol® DP0 and DP30 scaffolds are shown in **Figure 1**. Fibre diameter (13.0 ± 2.2 µm vs 12.7 ± 4.1 µm) and scaffold porosity (74.0 ± 3.2% vs 77.0 ± 2.7%) agreed reasonably well between DP0 and DP30. The fibre dispersion was lower (i.e. fibre alignment was higher) in DP0 scaffolds (9.6 ± 2.0°) compared to DP30 scaffolds (17.8 ± 4.6°, p < 0.05). Scaffolds of both materials exhibited some degree of fibre merging (**Figure 1** bottom row). Morphological scaffold data and dimensions of the scaffold samples for implantation are summarised in **Table 1**.

**Table 1.** Morphometric properties of the electrospun scaffolds

| Parameter | n | DegraPol® DP0 | n | DegraPol® DP30 |
| --- | --- | --- | --- | --- |
| Porosity | 7 | $74.0 \pm 3.2\%$ | 6 | $77.0 \pm 2.7\%$ |
| Fibre diameter | 3 | $13.0 \pm 2.2 \mu\text{m}$ | 3 | $12.7 \pm 4.1 \mu\text{m}$ |
| Dispersion (Goodness) | 6 | $9.6 \pm 2.0^\circ (0.97 \pm 0.01)$ | 6 | $17.8 \pm 4.6^\circ (0.98 \pm 0.01)$ |
| Width | 70 | $8.6 \pm 0.5 \text{ mm}$ | 68 | $8.1 \pm 0.5 \text{ mm}$ |
| Thickness | 70 | $1.2 \pm 0.2 \text{ mm}$ | 68 | $1.2 \pm 0.2 \text{ mm}$ |
| Length | 70 | $11.1 \pm 1.6 \text{ mm}$ | 68 | $11.1 \pm 1.5 \text{ mm}$ |

**Figure 1.**
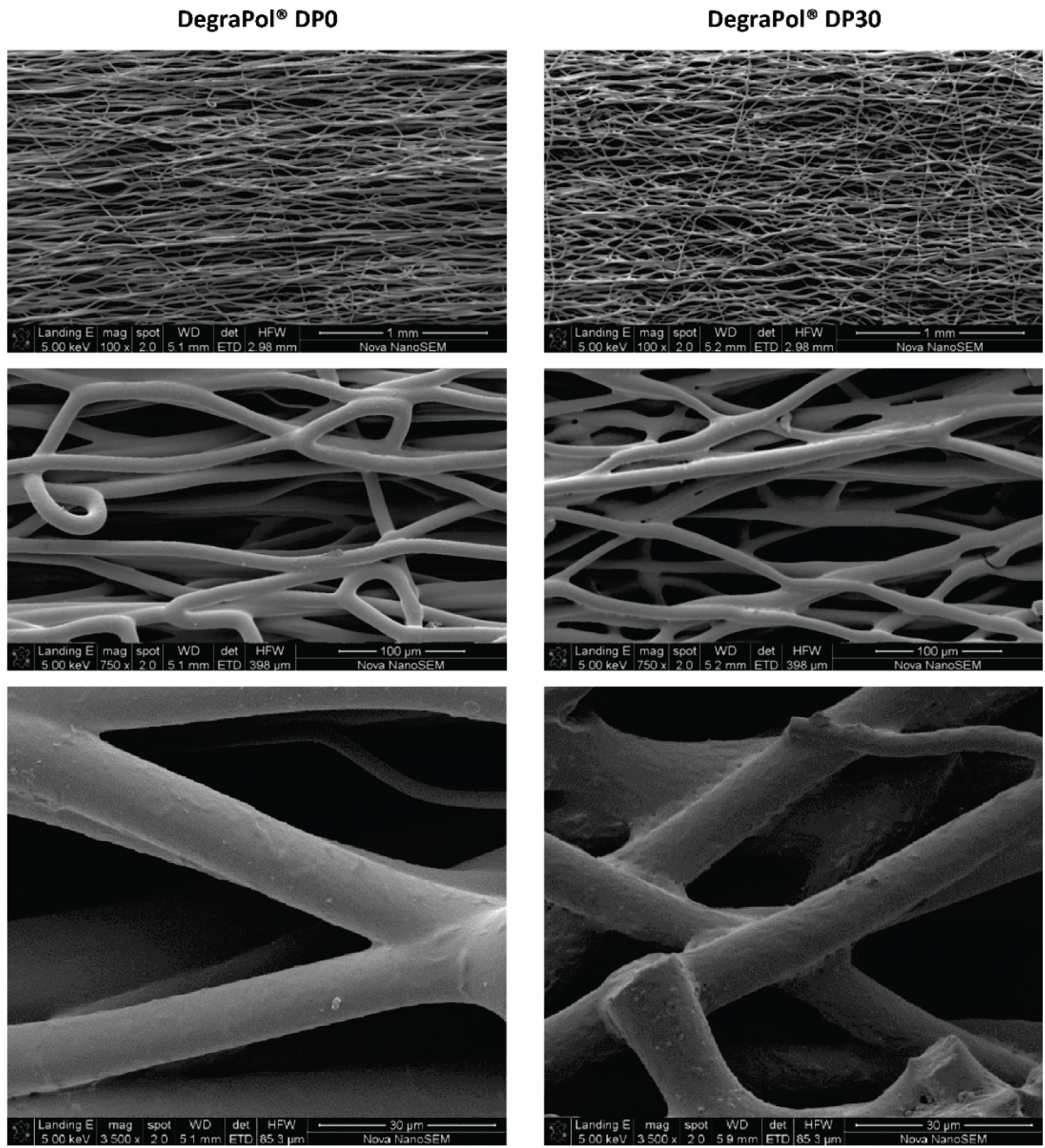
SEM micrographs of electrospun DegraPol® DP0 and DP30 scaffold at magnification of 100x (top), 750x (middle) and 3500x (bottom).

Weight-averaged molar weight of DegraPol® DP30 decreased significantly from 77.6 ± 5.7 kDa at T=0 days to 26.1 ± 6.7 kDa at T = 28 days, see **Figure 2**. Approximately half of this decrease occurred within the first 7 days (51%), whereas the decrease continued more gradually (T = 7 to 14 days: 19%, T = 14 to 21 days: 16%, T = 21 to 28 days: 14%). Biostable DegraPol® DP0 was analysed at T = 0 and 28 days only (n = 1) and did not exhibit a change in M_W_.

**Figure 2.**
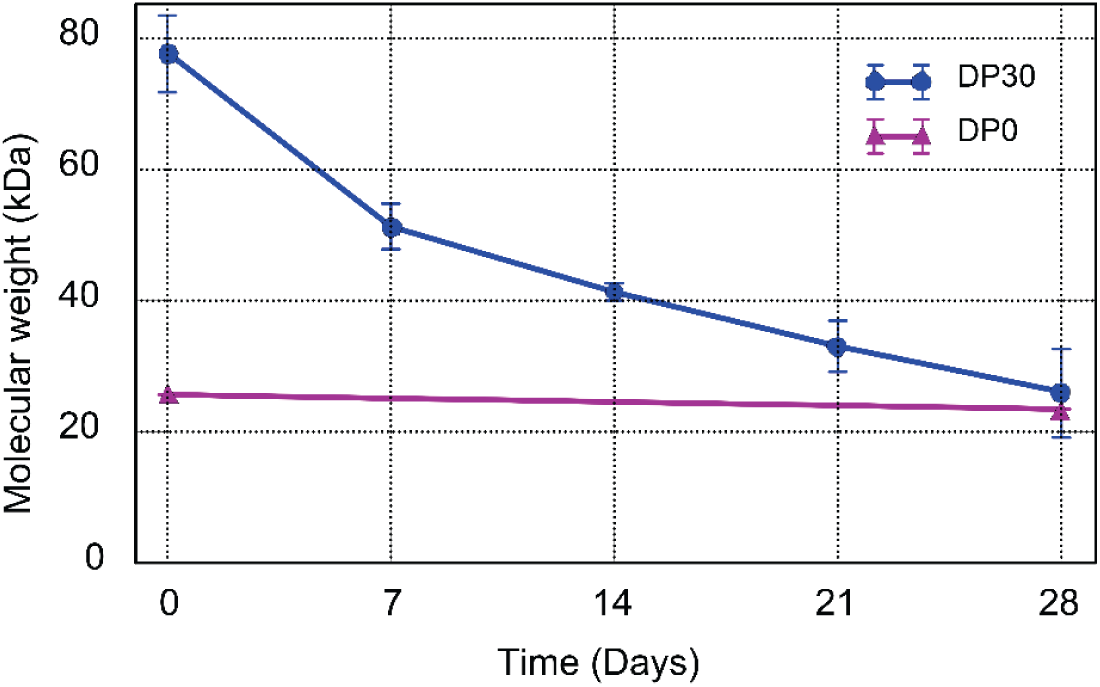
Molecular weight of DegraPol® DP0 and DP30 samples submerged in PBS versus exposure time up to 28 days.

Tissue ingrowth after 28 days was slightly higher, although not statistically significant, in DegraPol® DP0 scaffolds (83 ± 10%) compared to DegraPol® DP30 scaffolds (77 ± 4%). The progress of tissue ingrowth throughout the study is illustrated in representative micrographs of H&E sections of the scaffolds and quantified in **Figure 3**.

**Figure 3.**
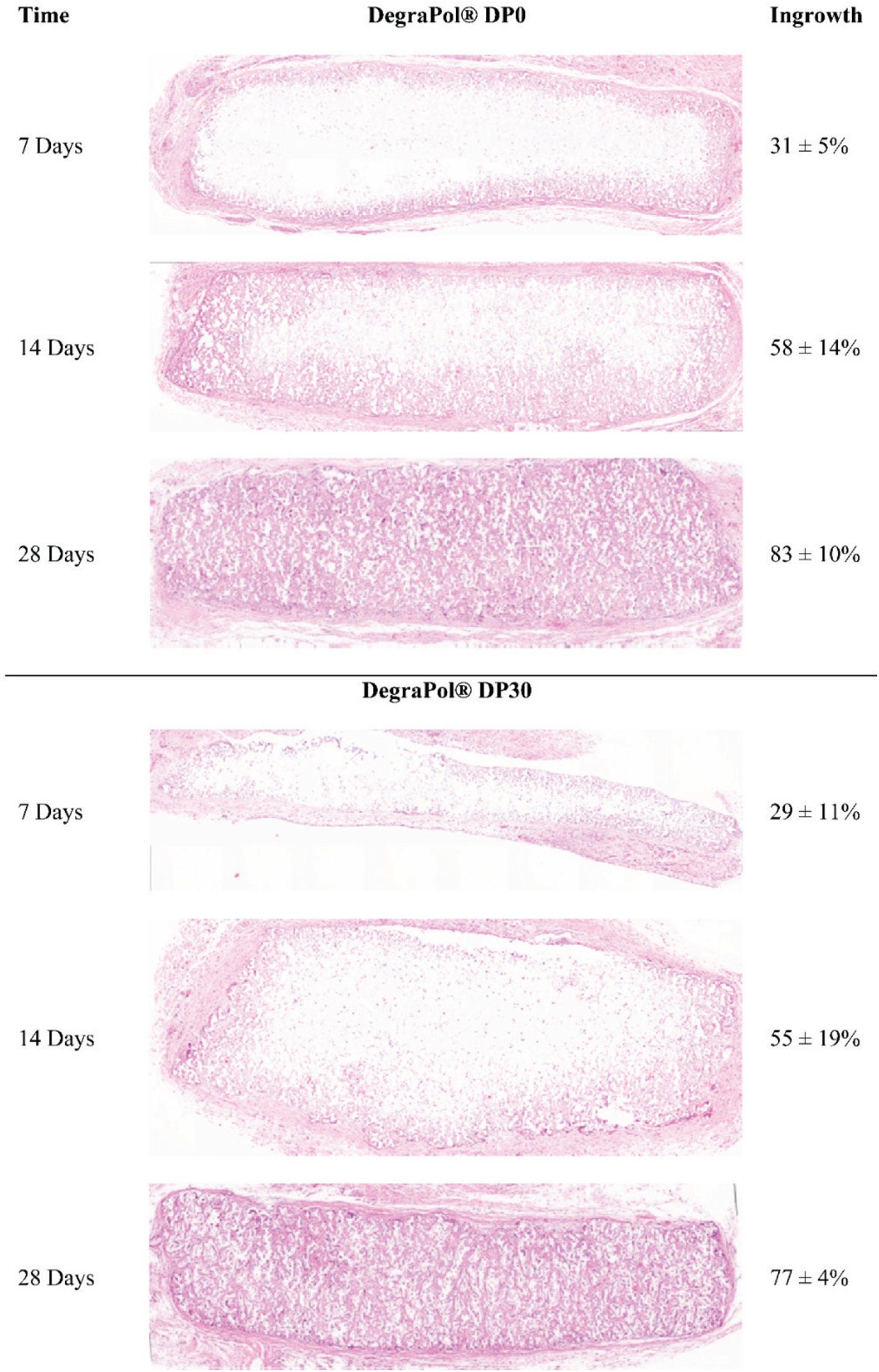
Histology micrographs of H&E sections in mid region of DegraPol® DP0 and DP30 scaffold samples and extent of tissue ingrowth after 7, 14 and 28 days of implantation.

**Figure 4 (a)** shows stress-strain data for axial and circumferential directions of pre-implant scaffold samples with and without load cycling. The resulting axial and circumferential stress scaling functions S_C_(ε) are illustrated in **Figure 4 (b)**. These functions quantify the impact of the load cycling for strains up to 16%. Disregarding data for strain below 2% (due to low signal-to-noise ratio), it was observed for axial DegraPol® DP0 scaffold samples that load cycling reduced the stress, σ_Cycled_, to 0.43% of the stress of non-cycled scaffolds, σ_Noncycled_, for strain, ε, between 2% and 12%. For ε ≥ 12%, load cycling decreased the stress to σ_Cycled_ = 0.33 σ_Noncycled_, indicating further weakening of the scaffold. For circumferential DegraPol® DP0 samples, σ_Cycled_ gradually increased from 0.20 σ_Noncycled_ at ε = 2% to 0.59 σ_Noncycled_ at ε = 12% strain, and decreased to 047 σ_Noncycled_ at ε = 16%. For DegraPol® DP30 scaffold, load cycling resulted in stress between σ_Cycled_ = 0.80 σ_Noncycled_ at ε = 2% and σ_Cycled_, = 0.98 σ_Noncycled_ at ε = 12% and 16% in axial scaffold direction, compared to σ_Cycled_ = 0.26 σ_Noncycled_ at ε = 2%, 0.93 σ_Noncycled_ at ε = 12%, and 0.97 σ_Noncycled_ at ε = 16% in circumferential scaffold direction.

**Figure 4.**
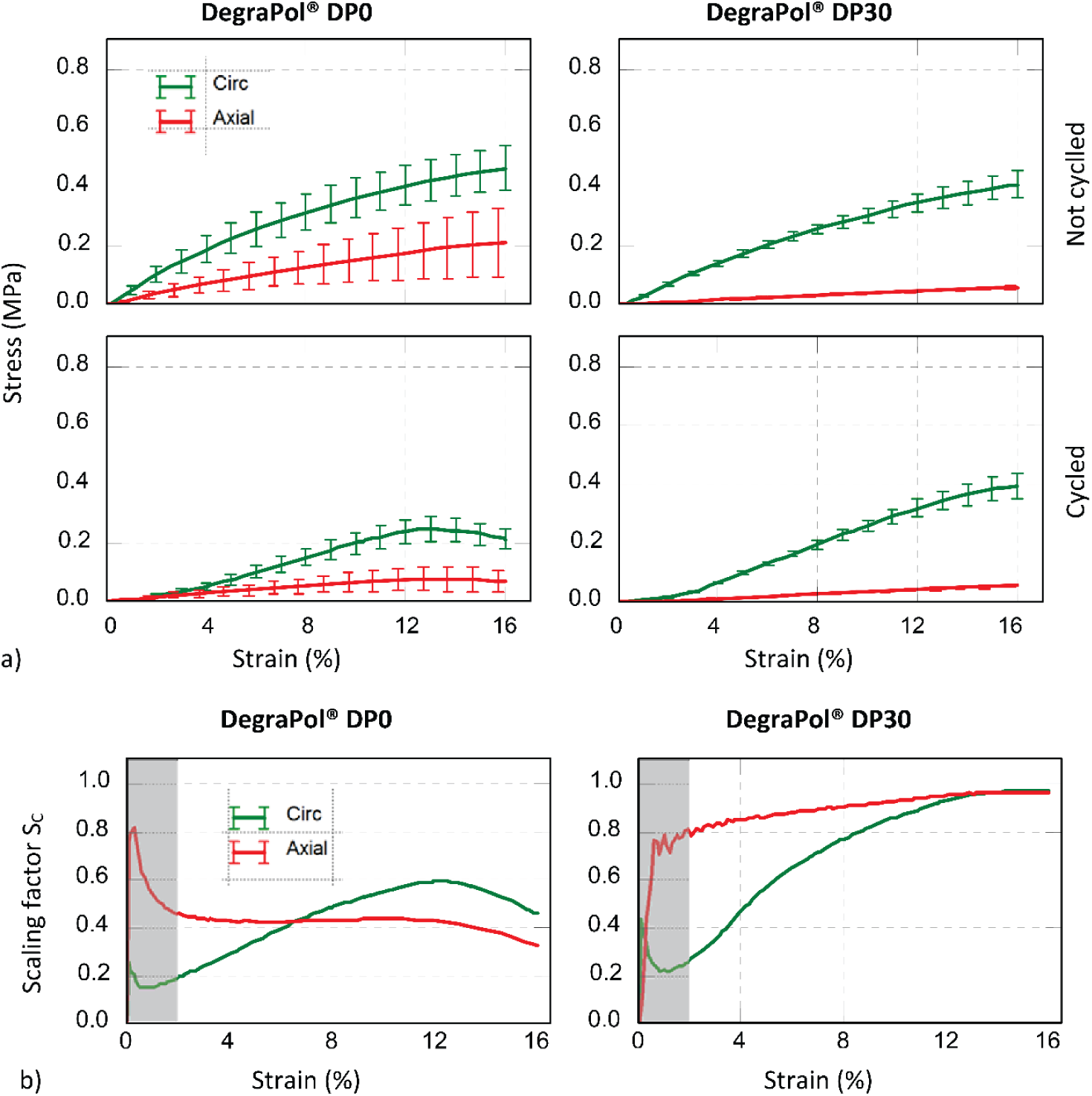
a) Stress versus strain curves for circumferential and axial direction of DegraPol ® DP0 and DP30 scaffolds before implantation (T = 0 day) for tensile loading protocol without and with load cycling; b) Stress scaling factor S_C_ = σ_Cycled_/σ_Noncycled_ versus strain, obtained from data shown in (a), representing the effect of load cycling on stress-strain behaviour of DegraPol ® DP0 and DP30. Data for strains below 2% were not considered due to high signal-to-noise ratio of force data based on low force magnitudes.

**Figure 5** illustrates stress-strain data obtained without load cycling as measured for DegraPol® DP0 and DP30 scaffolds in circumferential and axial directions at the various time points of the study. Stress-strain data after incorporation of the effect of load cycling using the stress scaling function S_C_(ε) are provided in **Figure 6**; the mechanical characteristics σ_12%_, σ_16%_, E_6%_ and E_12%_ are shown in **Figure 7**. The changes of stress and elastic modulus from 0 to 14 and from 14 to 28 days are summarized in **Table 2**.

**Table 2.** Changes in stress and elastic modulus characteristics from 7 to 14 and 14 to 28 days for DegraPol® DP0 (p < 0.05 for all values).

| Parameter | Change from<br>T = 7 to 14 days | Change from<br>T = 14 to 28 days |
| --- | --- | --- |
| $\sigma_{12\%}$ | +67% | -57% |
| $\sigma_{16\%}$ | +78% | -49% |
| $E_{6\%}$ | +63% | -52% |
| $E_{12\%}$ | +87% | -46% |

**Figure 5.**
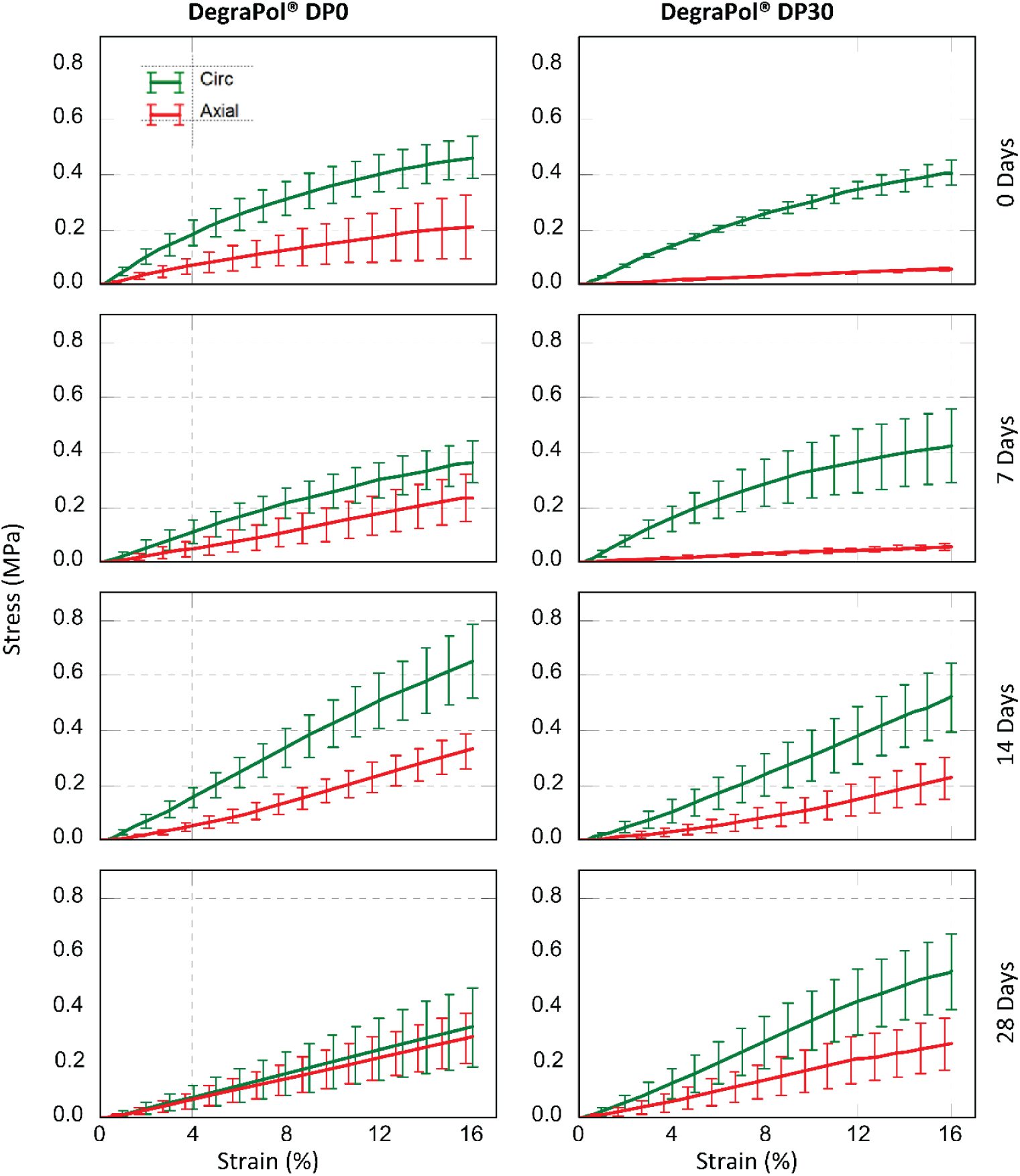
Stress versus strain curves for circumferential and axial direction of DegraPol ® DP0 and DP30 scaffolds without load cycling before implantation (0 day) and after implantation for T = 7, 14 and 28 days.

**Figure 6.**
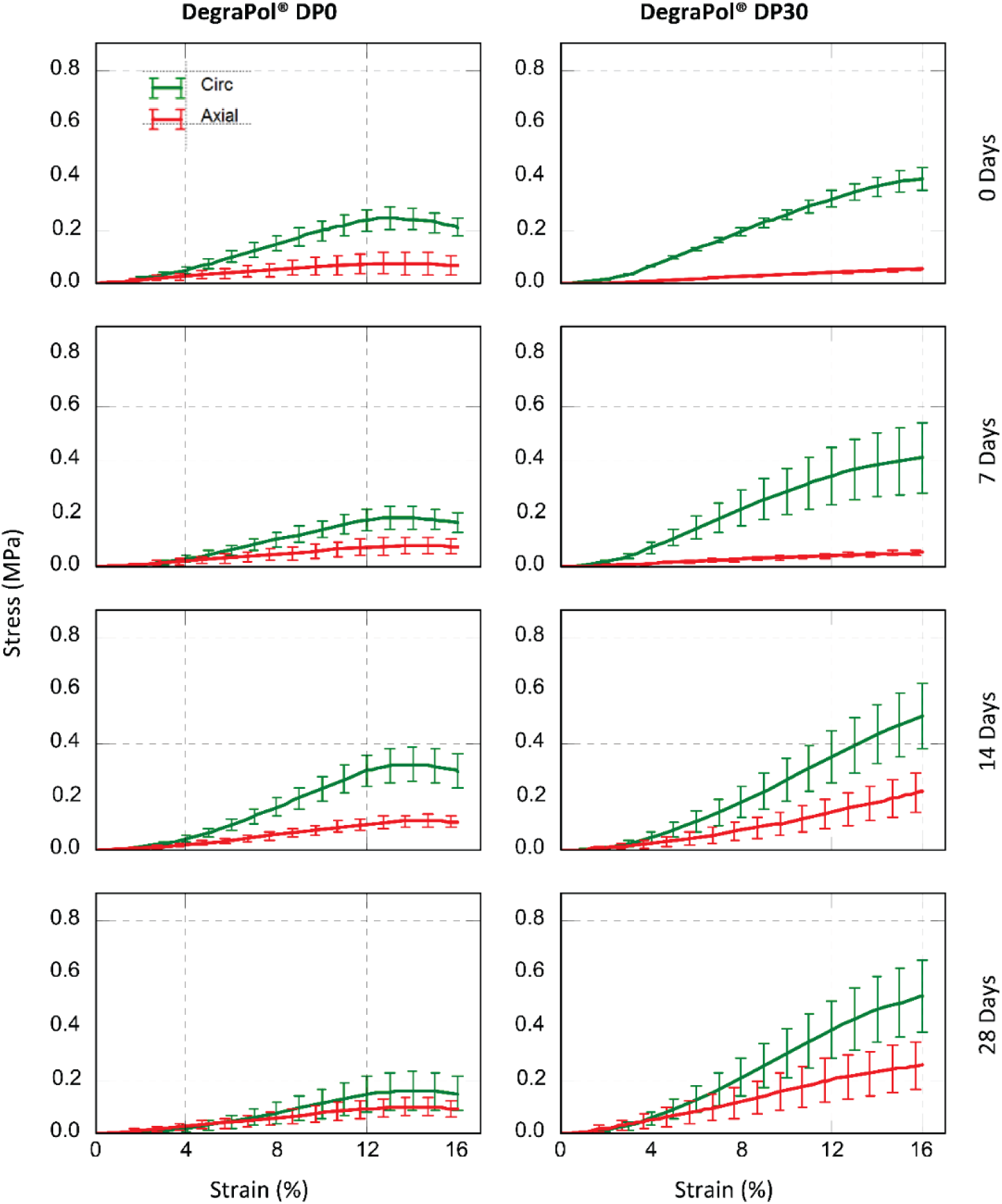
Stress adjusted for load cycling versus strain curves for circumferential and axial direction of DegraPol ® DP0 and DP30 scaffolds before implantation (0 day) and after implantation for T = 7, 14 and 28 days.

**Figure 7.**
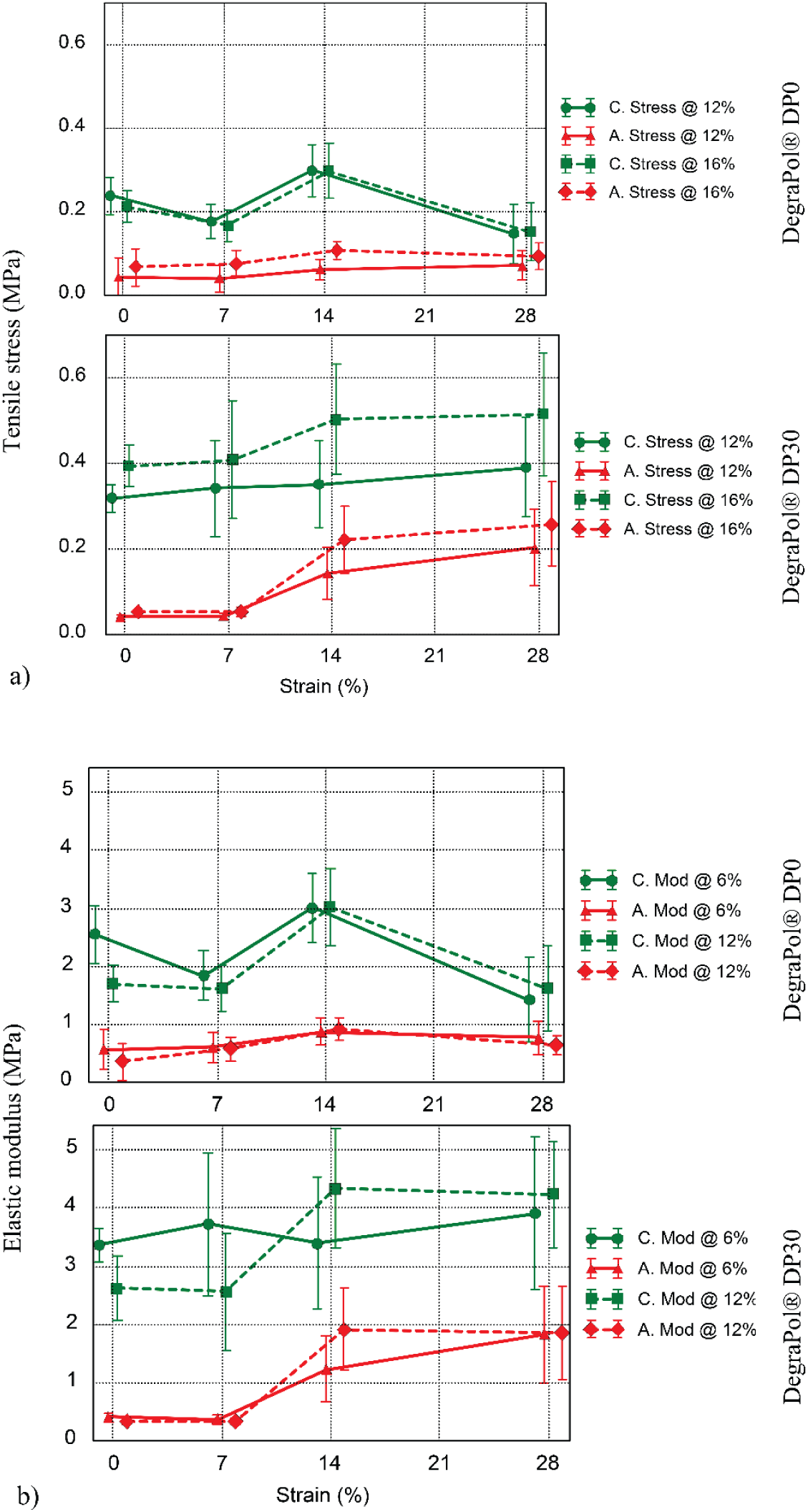
a) Stress at a strain of 12% and 16% versus implantation time and (b) elastic modulus at a strain of 6% and 12% versus implantation time for the circumferential (C) and axial (A) direction of DegraPol® DP0 and DP30 scaffold implants.

## 4. Discussion

The current study facilitated the paired *in vivo* assessment of the mechanics of biostable and degradable electrospun scaffolds based on the same polyester-urethane formulation to isolate the effects of material degradation and tissue formation after implantation. Biostable and biodegradable polyester-urethane scaffolds were implanted in the same animal.

The mechanical properties of biostable DegraPol® DP0 and degradable DegraPol® DP30 were found to be in the same range, although different responses were observed over implantation time. For DegraPol® DP0, circumferential data, but not axial, showed a significant increase from T = 7 to 14 days, and a significant decrease from T = 14 to 28 days in all characteristic measures, namely stress at 12% and 16% strain, and elastic modulus at 6% and 12% strain (**Table 2**). DegraPol® DP30 scaffolds exhibited a significant increase in stiffness *in vivo*. Whereas E_6%_ did not change, E_12%_ increased between T = 7 to 14 days by 50% (p < 0.05) for circumferential samples and by 213% for axial samples. This increase in elastic modulus was not reflected in the stress of circumferential samples but was for axial samples. Stress at 16% remained constant between T= 0 and 7 days but increased by 296% at T = 14 days (p < 0.05) and remained at that level at T = 28 days. The increase in elastic modulus after T = 7 days was linked to tissue ingrowth. Although there was approximately 30% ingrowth by 7 days, it was not yet fully interconnected throughout the scaffold, and thus did not contribute much to the overall mechanics. As tissue continued to populate the scaffold, regions with neo-tissue interconnected and developed greater structural strength.

Consistent mechanical performance from implantation through healing and scaffold degradation is highly desired for tissue-regenerative scaffolds, e.g. when used for vascular grafts. In the early phase after implantation, the circumferential stiffness in the scaffolds is based predominantly on fibres. As scaffold fibres degrade, the scaffold stiffness will decrease, unless the tissue balances out the difference. The consistency observed in circumferential stiffness of DegraPol® DP30 scaffold was thus of specific importance. Our previous *in vitro* study showed that the loss in material strength of DegraPol® DP30 predominantly occurred in the first 14 days [17], which correlated with decrease in molecular weight by T = 14 days. Assuming a similar trend for the scaffold *in vivo*, it holds that the tissue incorporation was contributing to the overall stiffness and balancing out the loss due to degradation. The stability in circumferential mechanical response contrasts, however, with the results by Teutelink et al. [22] where the grafts not implanted were significantly stiffer than the neo-vessels explanted after 90 days.

DegraPol® DP0 and DP30 scaffolds both featured a substantial degree of fibre alignment. However, DegraPol® DP0 scaffolds showed higher fibre alignment than DegraPol® DP30 scaffolds (p < 0.05), with high goodness values for the Gaussian fits of the fibre dispersion data in both cases (**Table 1**).

Tissue ingrowth was similar in DegraPol® DP0 and DP30 scaffolds, although it appeared to be slightly slower in the latter (**Figure 3**). Levels of tissue ingrowth increased significantly between all subsequent time points of the study for both DegraPol® DP0 and DP30 scaffolds (p < 0.05) except the increase between T = 14 and 28 days for DegraPol® DP30.

In order to avoid inducing premature damage in DegraPol® scaffold samples at later time points of the study, load cycling was not included in the protocol for mechanical characterisation. However, the effect of load cycling was characterised for pre-implant scaffolds and applied to scaffolds retrieved at the various time points. DegraPol® DP0 scaffolds displayed a higher sensitivity to load cycling than the DP30 scaffolds. For DP0 scaffolds, the stress scaling parameter SC ranging between 0.20 and 0.59 indicates that load cycling reduced stress in the scaffold by at least 41%. In addition, the decrease in stress σ_Noncycled_ (**Figure 4(a)** bottom left panel) and stress scaling parameter S_C_ (**Figure 4(b)** left panel) for strain above 12% indicates damage induced in the DegraPol® DP0 scaffold due to load cycling) both in direction of and transverse to the fibre alignment. For DegraPol® DP30, S_C_ ranged between 0.26 and 0.98 across the strain range studied. The DegraPol® DP0 scaffolds showed a more substantial weakening due to pre-cycling when compared to the DegraPol® DP30 scaffolds.

DegraPol® DP0 scaffolds were significantly more aligned than DegraPol® DP30, with a mean dispersion angle of less than 10°, compared to >17° for DegraPol® DP30. DegraPol® DP30 scaffolds also showed more fibre merging than DegraPol® DP0, another possible factor. Lee et al. [23] reported similar behaviour in Pellethane® electrospun meshes. They indicated that a higher degree of fibre merging resulted in better distribution of load compared to an otherwise similar structure with lesser fibre merging. Low degrees of fibre merging led to a structure in which the load was not as readily distributed among fibres, resulting in earlier onset of failure of individual fibres, and consequently the entire mesh. This correlated with the observation that DegraPol® DP30 fibres displayed more merging and higher tolerance to cyclic loading than the DegraPol® DP0 scaffolds. In addition, higher alignment led to individual fibres experiencing higher strains at low scaffold strain and thus failing earlier, as the fibres were already straight and thus did not need to first straightened out [24]. Again, the scaffold morphology correlated with tensile test results, with the higher aligned DegraPol® DP0 scaffolds incurring more damage at lower strains, as reflected by the decrease in stress post 12% strain.

## 5. Conclusions

Considering our previous *in vitro* observation that the DegraPol® DP30 scaffold loses its material strength predominantly during the first 14 days of hydrolytic degradation, the consistency of the elastic modulus of the degradable scaffold between 14 and 28 days in the current study is an indication that the regenerated tissue construct retains its mechanical properties. The strain stiffening of the tissue constructs with DegraPol® DP30 scaffold from 7 days of implantation onwards (evidenced through the increase of elastic modulus from 6% to 12% strain) is an additional advantage and mechanical behaviour observed in soft biological tissues such as blood vessels. For the application of the scaffold as tissue regenerative vascular grafts, further research is required to indicate whether similar results will be obtained in the circulatory implant position.

## Acknowledgements

The authors thank ab medica S.p.A for donating the DegraPol® material for this study. ETH Zurich and the University of Zurich are owners and ab medica S.p.A is exclusive licensee of all Intellectual Property Rights of DegraPol®.

## Data

Quantitative data for fibre diameter, fibre alignment, tissue ingrowth and mechanical characterisation are available on ZivaHUB (http://doi.org/10.25375/uct.9892367).

